# 3D super-resolution imaging using a generalized and scalable progressive refinement method on sparse recovery (PRIS)

**DOI:** 10.1101/532143

**Authors:** Xiyu Yi, Rafael Piestun, Shimon Weiss

## Abstract

Within the family of super-resolution (SR) fluorescence microscopy, single-molecule localization microscopies (PALM[1], STORM[2] and their derivatives) afford among the highest spatial resolution (approximately 5 to 10 nm), but often with moderate temporal resolution. The high spatial resolution relies on the adequate accumulation of precise localizations of bright fluorophores, which requires the bright fluorophores to possess a relatively low spatial density. Several methods have demonstrated localization at higher densities in both two dimensions (2D)[3, 4] and three dimensions (3D)[5-7]. Additionally, with further advancements, such as functional super-resolution[8, 9] and point spread function (PSF) engineering with[8-11] or without[12] multi-channel observations, extra information (spectra, dipole orientation) can be encoded and recovered at the single molecule level. However, such advancements are not fully extended for high-density localizations in 3D. In this work, we adopt sparse recovery using simple matrix/vector operations, and propose a systematic progressive refinement method (dubbed as PRIS) for 3D high-density reconstruction. Our method allows for localization reconstruction using experimental PSFs that include the spatial aberrations and fingerprint patterns of the PSFs[13]. We generalized the method for PSF engineering, multi-channel and multi-species observations using different forms of matrix concatenations. Reconstructions with both double-helix and astigmatic PSFs, for both single and biplane settings are demonstrated, together with the recovery capability for a mixture of two different color species.

## 1. INTRODUCTION

Super-resolution (SR) fluorescence microscopy is an indispensable tool for biological and biomedical research [14-17]. Within the family of optical SR technologies, localization based methods, such as PALM[1], STORM[2] and their derivatives, exhibit among the highest spatial resolution (approximately 5 to 10 nm), but at the disadvantage of lower temporal resolution. Higher time resolution requires a faster accumulation of localized fluorophores, which can be achieved with localizations at higher emitter densities that require less frames of camera acquisitions. Such methods include fittings of multiple emitters that are demonstrated to work with moderate to high densities [18, 19], and compressive sensing methods such as CSSTORM[3], L1-homotopy[4] and SOFI inspired sparse recovery [20, 21].

At the same time, advances in designing three dimensional PSFs, such as double-helix PSF[5, 11], astigmatic PSF[6], saddle-point PSF[22], and tetrapod PSF[23], has led to fast growing interests and applications of localization based SR microscopy for thicker samples [24]. At the single molecule level, Aristov et al developed ZOLA-3D[25] that has enabled flexible 3D localization over an adjustable axial range. At the higher density conditions: Barsic et al utilized either matching pursuit or convex optimization combined with a PSF dictionary and sparsity constraint to solve for the emitter localizations [5]. Junghong et al introduced sparse recovery followed by refinement of localizations (FALCON^3D^) and demonstrated its utility for an astigmatic/biplane imaging system [6]. Shuang et al developed a similar approach with open source software package that was validated on various 3D PSFs [7].

In addition, several further advances have been achieved at the single-molecule level with integrations of better fitting models, optics engineering and extra observation channels. For example, by fitting towards an experimental PSF model that compensates for most of the optical aberrations, minimal uncertainty of localization fitting have been achieved with video rate localizations[13]. By combining extra observation channel with modified optics, a mixture of multi-color species can be identified with SR[8, 9]. Similarly, using engineered PSFs that encodes the information of dipole orientation [26] or spectra information [12], the extra information can be encoded and recovered with SR.

However, such integrations are challenging in 3D under high-density conditions using the existing framework of open source implementation[7]. This is because the algorithm solves a sub-problem in the Fourier space, which indeed significantly reduces the computation cost, but causes the method to rely on the convolution assumption that requires the PSF to be translational invariant, therefore, without the ability to account for optical aberrations. We note here that when the PSF exhibits spatial variation, the observation is no longer a strict convolution between the ground truth and the PSF, therefore, the convolution assumption needs to be avoided. In order to extend the methodology advancements achieved at the single-molecule level [7-9, 12, 13, 26], a more generalized algorithm is needed for the recovery of 3D, high-density, multi-channel and multi-species conditions.

In this work, we introduce a systematic progressive refinement method for sparse recovery (PRIS) for 3D super-resolution microscopy, which is generalized and scalable for different imaging system with synchronization capability. PRIS keeps the form of simple matrix operations without relying on the convolution assumption of the observation process, allowing for convenient incorporation of experimental PSF, different background components, different species of signal sources and parallel measurements from different imaging channels. The sensing matrix and the reconstruction vector are defined with coarse discretization initially, and refined progressively and regionally to reach finer discretization. Such regional refinement enables higher resolution reconstruction without the drastic increase in the computational cost. In principal, the discretized location coordinates in the 3D space may also be extended into a hyper-dimensional space to include extra dimensions of information, or different PSF species that co-exist and overlap in the same measurement, such as dipole orientation [26], and color [12]. We have validated PRIS and the associated generalization strategy with realistic simulations for single-or multi-channel observations (biplane) and mixture of two color species, with 3D PSFs of either Double-Helix PSF (SPINDLE[27]) or astigmatic PSF.

The rest of this paper is organized as follows: In section 2, we introduce the theory and algorithm used in our work, including both review of dependencies and emphasis on our contribution. In section 3, we validate our method and the associated generalization strategy with simulations under various conditions. We also characterized the performance of our method. In section 4, we provide a discussion highlighting the advantages of our method.

## 2. THEORY AND ALGORITHM

### 2.1. L1-norm regularized sparse recovery with progressive refinement

The image formation process of fluorescence microscopy can be described by a linear mapping process. Mathematically, we have:

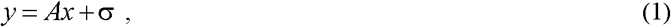

where ***x*** is a vector (we dub as the target vector) accounting for all possible signal sources, ***y*** is the observation vector corresponding to the observed image, ***A*** is the sensing matrix (observation matrix) describing the linear mapping from the signal source to the observation, and ***σ*** represents an additive noise component. The effect of omitting the Poisson noise is negligible as demonstrated by the existing compressive sensing applications for SR microscopy[3, 4, 20, 21]. The following L1-norm regularized sparse recovery solves for ***x*** when ***A*** and ***y*** are known and with the prior knowledge that ***x*** is sparse (possesses a small portion of non-zero entries):

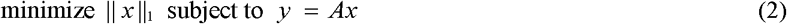

The observation vector ***y*** is obtained by vectorizing the observed (or simulated observation) image. An empty vector ***x*** is defined to represent a collection of location coordinates in a discretized manner, and the sensing matrix ***A*** is calculated using the knowledge of the observation process, for example, as provided by an experimentally characterized PSF or a theoretical PSF model. In cases when the noise component exhibits non-zero average, extra columns can be added to the ***A*** matrix to account for different background components[6], or an offset. In this work, we adopt the fast linearized Bregman iteration[28] that constrains the inverse problem in (2) as follows:

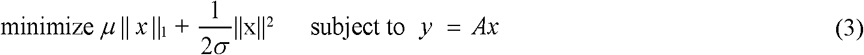

Although (3) is different from (2), it has been shown to be equivalent to (2) for large *μ* [29]. As shown in Algorithm 1, linearized Bregman iteration involves only simple matrix/vector operations and shrinkage[28] operation consisting only Boolean operation and subtractions.

#### Algorithm 1. Basic linearized bregman iteration

1. Calculate the sensing matrix ***A*** based on the candidate signal sources and the PSF model.

2. Obtain the ***y*** vector from the observation image.

3. Define ***μ*** and ***σ***, and initialize the iteration index ***k*** as 0.

4. Initialize ***x***_0_ and ***u***_0_ as zero vectors, with length equals to the total number of candidate signal sources.

6. While “‖ ***y*** – ***A·x***‖ not converge”, do:

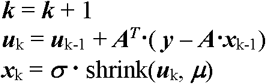

End while, recovered ***x = x***_***k***_

Note that shrink(.) is the shrinkage operator[28]. We have incorporated fast linearized Bregman iteration with “kicking” to improve the convergence speed. Interested readers are referred to [28] for more details.

To bring more insight, we discuss here the physical meaning of the formula of our sparse recovery task. In the basic form for 3D recovery, one element in the ***x*** vector (denote the *i*^th^ element as ***x***_***i***_) represents one candidate position for the emitters in the sample. The ***x*** vector corresponds to a group of voxels that collectively represent a 3D volume in a discretized manner, and the value of ***x***_*i*_ represents the total signal emitted from the corresponding voxel. In addition, extra elements can be added to ***x*** to account for extra possible signal sources, and the corresponding observation profile produced by each candidate needs to be added as extra columns in ***A***. With further generalization, one element in ***x*** could represent any candidate signal source defined by multiple dimensions of information in addition to the location coordinates, such as dipole orientation, spectra, etc. We note here that the *i*^th^ column in ***A*** represents the unit profile of the observation created by the *i*^th^ possible source, as corresponding to ***x***_*i*_. In other words, the multiplication of the *i*^th^ element in ***x*** and the *i*^th^ column in ***A*** yields a linear component (i.e. additive component) in the observation vector ***y***, while the value of ***x***_***i***_ is the coefficient of this linear component, and the *i*^th^ column in ***A*** is the unit observation profile created by the *i*^th^ signal source.

In order to push for optimum performance of 3D recovery of a thick sample, the inverse problem needs to account for a sufficiently large 3D space with high sampling rate. Therefore, the total number of voxels increases, leading to an increase of RAM requirement. For example: Assuming an input image patch of 64-by-64 pixels, with a cubic volume of 6.4×6.4×1 μm^3^ represented by close-packed voxels of size 16^3^ nm^3^, the RAM requirement for the sensing matrix ***A*** with single float precision is approximately 152.6 Gigabytes. We note here that setting up the problem with Fourier transforms[7] can significantly reduce the RAM requirement, but relies on the convolution assumption that we explicitly would like to avoid as discussed in the previous section.

The proposed progressive refinement method (PRIS) is designed as an iterative algorithm on top of the sparse recovery solver to address the computation cost, without assuming the formation of the observation image as a convolution process. As shown in Figure 1 and Algorithm 2, the PRIS iteration is initialized by constructing the initial inverse problem, where the candidate locations are represented by close-packed coarse voxels (Figure 1(a)), yielding ***A*** and an empty ***x*** vector to recover, and the ***y*** vector is obtained by vectorizing the observation image. We note here that a pre-processing step can be used to initialize the pool of candidates more compactly with even lower RAM requirement. For example, incorporation of a segmentation step as implemented in many existing tools for single molecule localization microscopy, but without discarding aggregates. Once the inverse problem is clearly defined, a sparse-recovery solver is used to solve for ***x***. The resultant ***x*** is inspected, and the non-zero elements in ***x*** are identified for the next PRIS iteration (Figure 1(c)). Specifically, the voxels corresponding to the non-zero ***x*** elements are refined into smaller voxels to yield a new set of candidate locations (Figure 1(d)), with which a new ***x*** and ***A*** will be constructed and combined with the original ***y*** to form a new sparse recovery task. The refinement process is repeated progressively (Figure 1(b)): Upon completion of each sparse recovery task, zero-value voxels are discarded, and non-zero voxels are refined to construct a new sparse recovery task for the next PRIS iteration. The sparse recovery solver inside the PRIS iterations is a modular component that is not restricted to our implementation. In Algorithm 2 we demonstrate the algorithm with linearized Bregman iteration in its basic form. In our implementation, we incorporated “kicking” [28].

**Figure 1.**
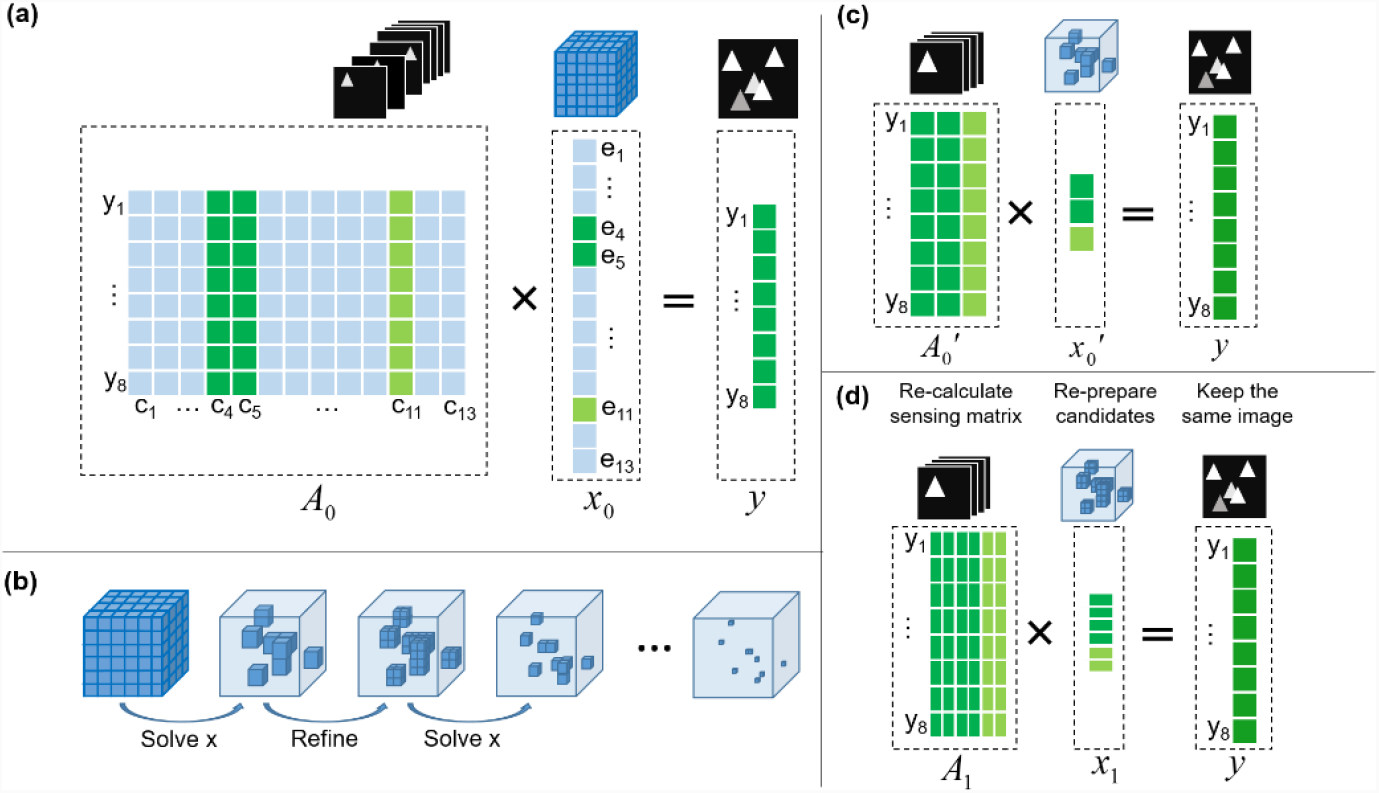
Sparse recovery with progressive refinement. (a) shows the formulation of the recovery problem ***Ax*** = ***y***, with known ***A*** and ***y***, to recovery ***x***. As labeled in (a), we highlighted the one-to-one correspondence between columns in ***A***, elements in ***x***, and the linear component in ***y***. (b)-(d) shows the progressive refinement process of sparse recovery.

#### Algorithm 2. Progressive Refinement method for Sparse recovery (PRIS)

1. Obtain the ***y*** vector (from the observation image); and obtain the system response function (i.e. **PSF**).

2. Define total PRIS iteration number **N**, and the list of PRIS refinement folds: **f** = {**f**_1_, **f** _2_, …, **f** _N-1_}. Typically, we have **N** = 4 or 5, and **f** = {1, 2, 2, …, 2}.

3. Initialize sample volume ***V*_0_**, discretization size ***D***_0_.

4. *<u>PRIS iteration begins:</u>* For PRIS iterations ***i*** = 0 to **N**-1:

4.1. If ***i*** > 0: Refine sample volume and discretization: ***V***_***i+1***_ = Refine(***V***_*i*_, ***x***_*k*_), ***D***_***i+1***_ = ***D***_***i***_/**f**_*i*_

4.2. Obtain the collection of candidate signal sources: ***C***_*i*_ = Sampling(***V***_*i*_, ***D***_*i*_).

4.3. Calculate sensing matrix ***A***_*i*_ based on ***C***_*i*_ and PSF model: ***A***_*i*_ = Sensing(***C***_*i*_, **PSF**).

4.4. Define ***μ*** and ***σ***, and set iteration index ***k*** = 0.

4.5. Initialize ***x***_0_ = 0, ***u***_0_ = 0 with lengths equal to the size of **C**_*i*_.

4.6. While “‖ ***y*** – ***A·x***_k_‖ not converge”:

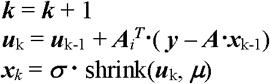

End while, recovered ***x = x***_***k***_

*<u>PRIS iteration ends.</u>*

5. Post processing and final result rendering using ***C***_*N-1*_ and recovered ***x***.

In summary, PRIS does not rely on the convolution assumption but can perform sparse recovery over a finely discretized pool of candidate signal sources. The reduction of RAM requirement is realized by progressively focusing the computation power onto smaller and regional subsections of the pool, allowing for recovery with larger volume and finer discretization.

### 2.2. Generalization of PRIS with matrix concatenations

With the spared RAM requirement, the advances achieved at the single molecule level can be easily integrated with PRIS using different forms of matrix concatenations. First, vertical concatenation of the sensing matrixes allows for synchronization of multi-channel observations, such as multiple focal planes, or different phase masks. Second, horizontal concatenation of the sensing matrixes allows for incorporation of multi-species of the signal source, such as background components[6], or different PSFs (such as different colors [12] or dipole orientation [26], etc.). Hybrid of horizontal and vertical concatenations can be utilized as well. In the general sense, PRIS can be applied to sparse recovery even at a hyperdimensional space[30], and the progressive refinement can be applied at each dimension independently. We explain below the vertical and horizontal concatenations of the sensing matrix, and further generalization can be realized recursively in the same manner.

As shown in Figure 2, the sensing matrixes are concatenated vertically, and the two different observation vectors are concatenated accordingly. The ***x*** vector remains the same, because the observations for both channels are generated by the same signal sources. Consequently, the overall sparse recovery task is reduced to the same form:

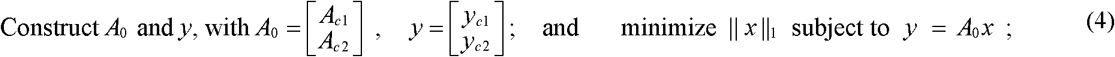

where ***A***_***c1***_ and ***A***_***c2***_ are the sensing matrixes corresponding to two different observation processes, ***y***_***c1***_ and ***y***_***c2***_ are the two different observations, and ***x*** is the recovery target. Such way of generalization can be applied to the recovery of different observations generated by identical signal sources (as circled by red dashed line in Figure 2 at the observation channels). Multiple observation channels could be, for example, biplane, or multiplane observations, etc. Accompanying information can be synchronized directly in the data analysis step via combined reconstruction. If the sample contains different species of signal sources that respond selectively to the observation channels (as labeled by cyan stars in Figure 2), then the species variation is underestimated by this strategy and a different generalization strategy for PRIS is required (discussed later).

**Figure 2.**
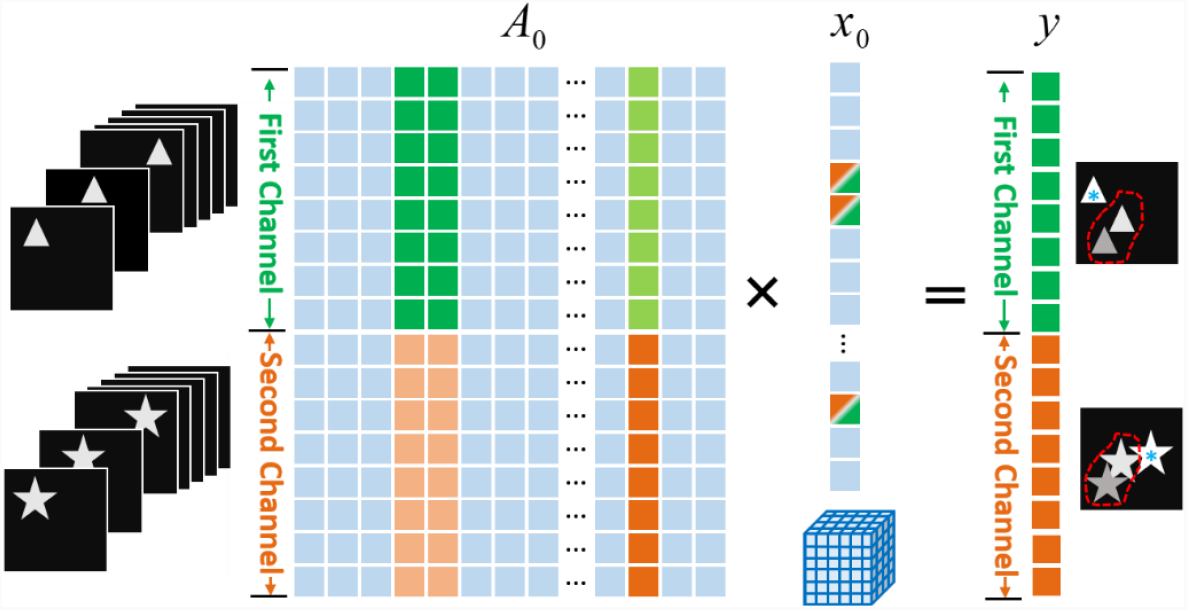
Generalization of PRIS with vertical matrix concatenations. Only the initial sparse recovery is illustrated, and the iteration with PRIS refinement is the same as shown in Figure 1. ***A***_***0***_ is the overall sensing matrix, and the concatenation is between two different sensing matrixes that describe the observations of two different channels. ***x***_***0***_ represents the pool of candidate signal sources that uniform for both observation channels, and ***y*** represents the overall observation as the combination of the separate observations from different channels.

Here we first discuss a generalization strategy for the case of two signal species observed through a single channel. As shown in Figure 3, the candidate locations for both species are initialized independently, and the corresponding voxels could be overlapping because both species exist in the same sample space. However, as PRIS iteration proceeds, the recovered candidate signal sources for each species can be located at different sets of voxels. Therefore, the refinement is performed independently for each species. In addition, we applied a soft species exclusion constraint to the fast linearized Bregman iteration solver. To be specific, as shown in Algorithm 2, the increment of ***μ***_k_ is ***A***_***i***_ **^*T*^*·***(***y*** – ***A·x***_k-1_), and we call it the increment vector. Similar to ***x***, the elements in the increment vector has a one-to-one correspondence to the voxels co-specified by the species tag and the 3D coordinates. In each iteration, for each doubly occupied voxel (defined as two voxels with the same 3D coordinates but different species tags), the corresponding two elements in the increment vector are inspected, and if the values of these two elements have the same signs, the element with a smaller amplitude is set to zero.

**Figure 3.**
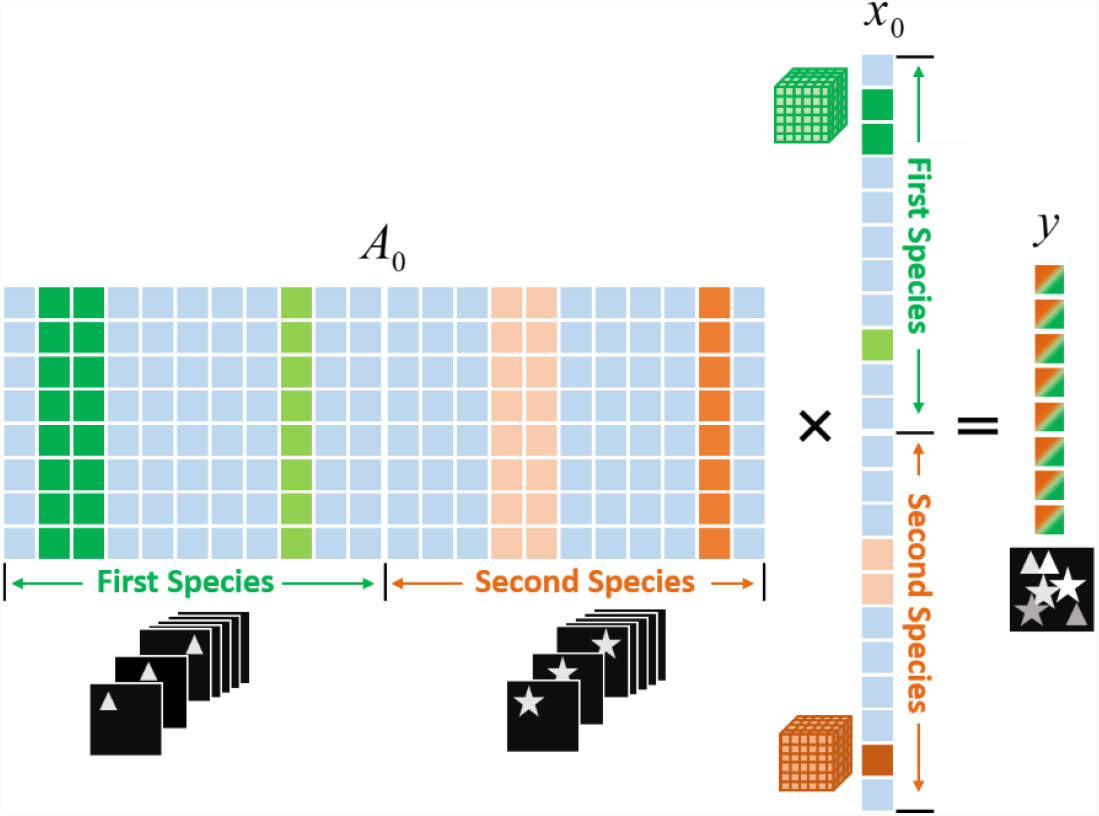
Generalization of PRIS with horizontal matrix concatenations. Only the initial sparse recovery task is illustrated and the iteration with PRIS refinement is the same as shown in Figure 1. ***A***_***0***_ is the overall sensing matrix that is the concatenation of two different sensing matrixes, each describing one portion of the observation contributed from an identical species through the same observation channel. ***x***_***0***_ represents the concatenated pool of candidate signal sources of two different species, and ***y*** represents the single channel observation with co-existence of two different species.

As shown in Figure 3, the sensing matrixes characterizing different species are horizontally concatenated that allows for the synchronization of different species. And the associated sparse recovery task has the conserved form as follows:

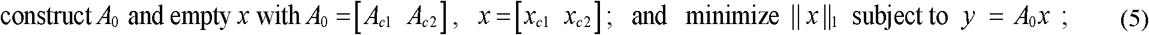

Following the similar strategy, hybrid concatenations can be constructed as well to account for different species observed through different channels, in case if a species produces no signal in part of the observation channels (as labeled by cyan stars in Figure 2), the corresponding sensing matrix can be a zero matrix. We remark here that the refinement has been previously used to improve the resolution and reduce the RAM requirement of sparse recovery for SR microscopy[6]. The merit of this work is the systematic progressive refinement and the generalization strategy for sparse recovery, while excluding the convolution assumption on the image observation process. The access to the fine discretization further allows for the classification step to identify the localization coordinates and efficiently reject dim localizations that could be false and misleading. Such merit allows for extension of a variety of different fluorescence imaging modalities and objects achieved at the single molecule level to the 3D high-density conditions, the increased time resolution of such methods through the faster accumulations of localizations can open up diversified possibilities for the study of dynamics and longer-range processes in complex systems. In the more general sense, PRIS can be generalized to sparse recovery at a hyperdimensional space with tensor formulation[30], and the progressive refinement can be applied at each dimension (each mode of the tensor) independently or collectively.

## 3. VALIDATION WITH SIMULATIONS

The performance of PRIS is validated and quantified on three set of simulations, and the general applicability of the method is demonstrated with two different 3D PSFs: astigmatic and SPINDLE PSF (a version of double-helix PSF) [27]. Details are explained below.

### 3.1. Recovery with astigmatic and SPINDLE PSFs with single plane observation

In this simulation, single plane observations are used with either SPINDLE PSF or astigmatic PSF to validate the performance of PRIS. A series of simulations are generated with randomly placed emitters in a 3D volume with a various total number of emitters. Figure 4 shows a representative recovery result for both PSFs. The PRIS result represents each candidate localization as a group of non-zero voxels. The recovered 3D voxels are classified into groups using the density based classification method[31], and further converted into a list of localization results for which the coordinates are calculated as the mass center of the voxels belonging to the same group, and the brightnesses information are obtained by taking the integral of the values of the voxels within the same group. The full set of simulated samples is used for further quantification, with a range of 10 to 400 emitters in a 3D volume of 6.4 μm in the lateral-(XY-) dimensions, and 1 μm in the axial-(Z-) dimension. The resultant simulations include different emitter densities ranging from 0.244 to 9.766 μm^−2^.

**Figure 4.**
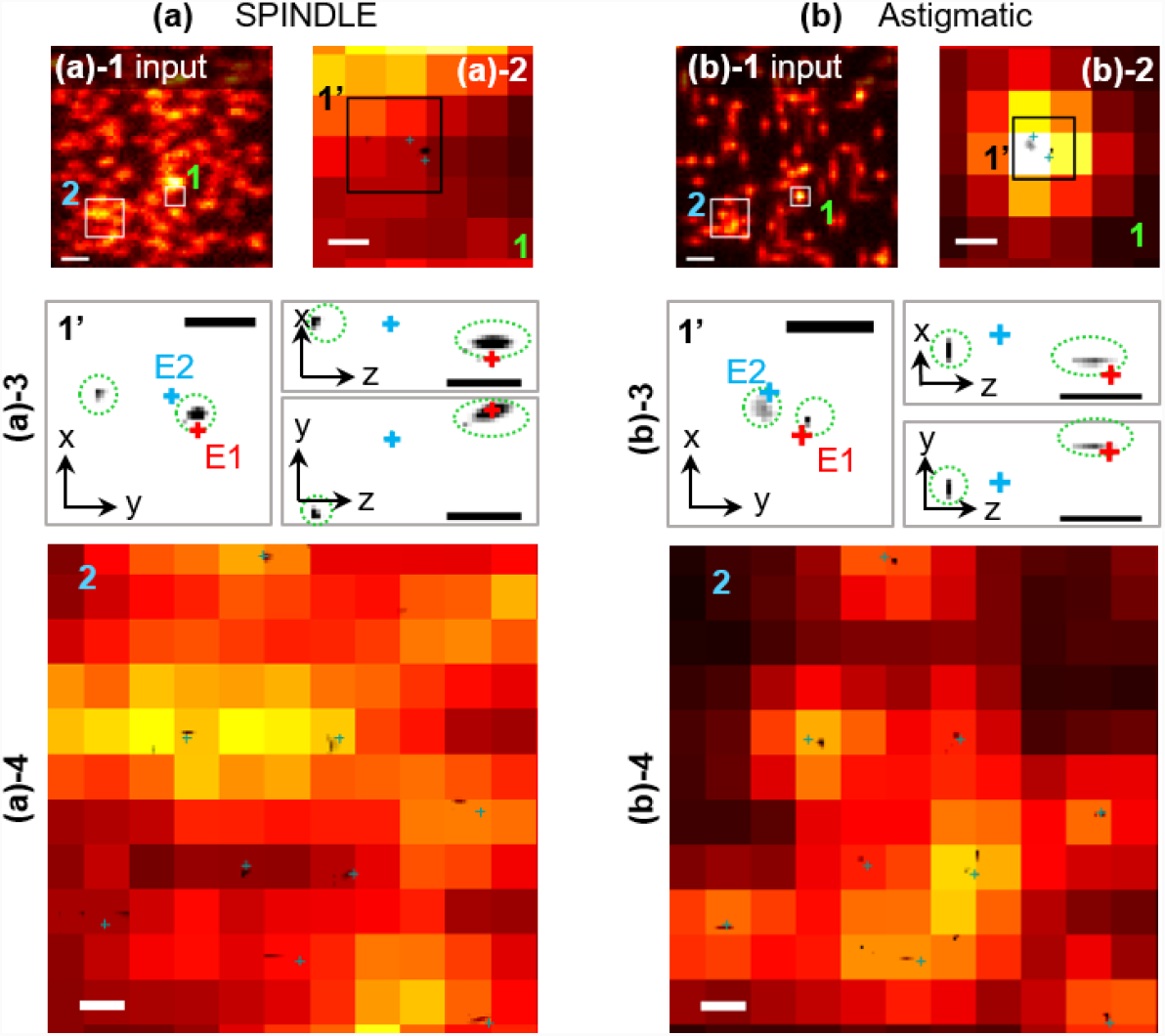
Representative recovery result for PRIS with (a) SPINDLE PSF and (b) Astigmatic PSF are shown here. (a/b)-1 are the input noisy blur images, where two zoom-in regions are labeled (as 1 and 2) and displayed in the rest of the panels. (a/b)- 2 shows the zoom-in region 1 with two emitters, which are further amplified into region 1’, as shown in (a/b)-3 for views in xy/xz/yz-planes. The reconstructed result is displayed at gray scale, appearing as groups of dark pixels (highlighted with circled with green dashed lines), and the ground truth emitter locations are labeled as red/blue crosses for E1 and E2 respectively. The red cross (E1) indicates a representative precise localization for both PSF models, with displacement of 23 nm for SPINDLE PSF, and 28 nm for Astigmatic PSF. Accordingly, the blue cross (E2) indicates a representative localization with worse precision for both PSF models, with displacement of 148 nm for double-helix PSF, and 66 nm for astigmatic PSF. The second region (Region 2, as labeled in (a/b)-1) represent a higher emitter density (equivalent to approximately 7.4 emitters μm^−2^), the corresponding zoom-in panels are shown in (a/b)-4. Scale bars in (a/b)-1 are 800 nm, scale bars in (a/b)-2/3/4 are 100 nm.

The photon budget for each emitter is 5000, and a background level of 100 photons are added to each pixel area. The simulated observation images are generated on an initial discretization of 10 nm pixels followed by 10×10 binning to reach a final pixel size of 100 nm. Photon counting noise (shot noise) is simulated with Poisson statistics. For each density condition, 20 simulations are performed with randomized 3D locations. We have quantified the performance of our method gauged by the recovered emitter density and localization precisions as shown in Figure 5. For the convenience of reading, we have plotted similar quantifications from previous works on the same graphs. We note here that the quantifications of different methods were not performed under the same simulation condition (as noted in Appendix, Table 1.). Our simulations represent among the challenging conditions. The SPINDLE PSF we have used is a non-optimized version of double-helix PSF. when comparing to the 2D cases, the signal from a single emitter expands over a larger area because of the larger PSF, and our signal is integrated over a smaller pixel area, resulting in a reduced SNR as compared to a more compact PSF and/or with larger integration area of the larger pixels.

**Figure 5.**
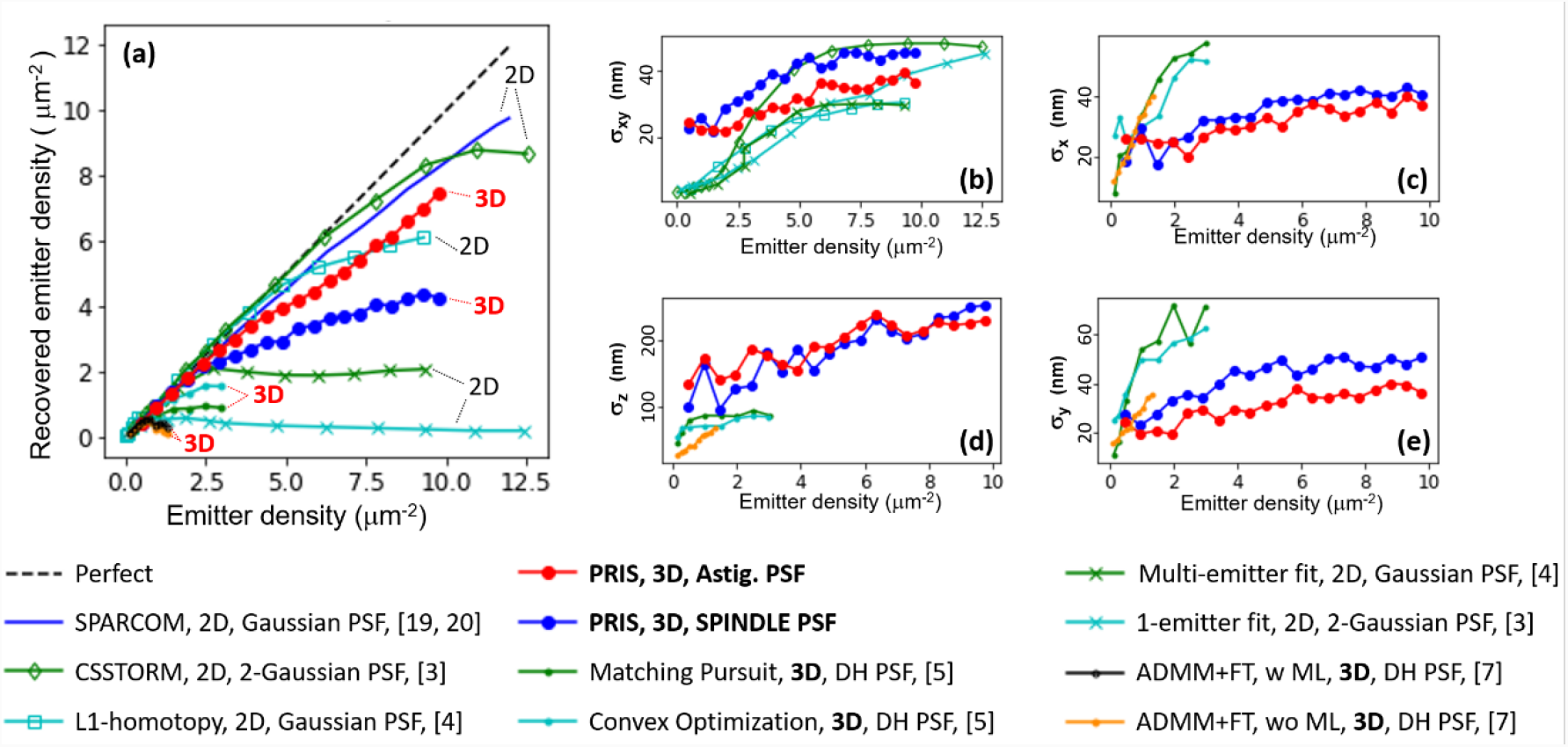
Performance of RIS recovery and previous works. (a) shows the recovered emitter density plotted against the ground truth emitter density. (b)-(e) shows the localization precision in the XY-, X-, Z-, Y- planes as a function of emitter density respectively. We can see that our method exhibits excellent 3D recovery capability with high fidelity at high-density conditions.

A simulated emitter is identified as recovered if a localization result is found within 1-pixel distance in the XY-plane. The densities of the recovered emitters are plotted against the actual emitter density (ground truth) in Figure 5(a). We can see that PRIS demonstrates excellent performance as compared to previous 3D recovery methods using compressive sensing [5, 7], and single- or multi-emitter fitting methods. We attribute such performance to the access of a very fine discretization (8.25 nm) granted by the progressively refined voxels. For the same reason, we have access to the much higher density conditions in our characterization. PRIS also exhibit comparable performance as compared to 2D approaches[3, 4, 20], regardless of the fact that the 3D PSFs used in our characterizations are imposing an intrinsically more challenging recovery task as compared to the 2D approaches where the regular PSFs are much more compact. When the PSF expands a larger area (SPINDLE), we expect an increase of the overlapping region, and the total budget of photons would also spread over a larger area, resulting in a lower SNR. The size difference between the astigmatic PSF and SPINDLE PSF can also explain the different performances of the two PSFs in our characterization.

The comparison of localization errors is performed after identification of correct localizations: A localization result is identified as correct if there exist a ground truth emitter within 1-pixel distance in the XY-plane. For all the correct localizations, the fitting error is defined as the displacement of the localization result from the ground truth, and the fitting precision is calculated as the standard deviation of the fitting errors. Figure 5(b) shows the comparison of PRIS with the 2D methods [3, 4, 20] by comparing the standard deviation of fitting errors in the XY-plane (dubbed as σ_xy_). We can see that PRIS demonstrate comparable or better performance in the lateral precision at high-density conditions (> 2.5 μm^−2^). Precision comparison with 3D methods [5, 7] are shown in Figure 5(c) to (e), where PRIS excels under high-density conditions (> 1 μm^−2^) among all the 3D methods [5, 7]. We note here that a least square fitting approach was applied to the localization result to finalize the localization result in the previous 3D approach[7], which is not implemented in our work, but is compatible with our method and is expected to further improve the performance of PRIS.

### 3.2. Recovery with multi-channel observation and single species (vertical concatenation)

Further simulations are performed to validate the generalization scheme of our method that utilizes vertical or horizontal concatenation of the sensing matrixes. Simulation of biplane observations of a pure color species is used to validate the vertical concatenation strategy, and single-channel observation for a mixture of two species with two different colors is used to validate the horizontal concatenation strategy.

Figure 6 shows the results for dual-channel observation simulations. The simulated sample contains 100 emitters randomly positioned in the sample volume of size 6.4×6.4×1 μm^3^. Different observations are generated to simulate biplane and single-plane observations with either SPINDLE and astigmatic PSFs, while the photon budget is maintained at 5000 counts per emitter. Figure 6(a) shows the single plane observation for both SPINDLE and astigmatic PSF separately, and Figure 6(b) shows the corresponding biplane observation for both PSF models separately. Due to the split of the dual planes, the total photons also split between the two observation channels (accounted in the simulations). The SNR difference between the observation images for single and biplane observations is due to this split. The recovery is performed through the vertical concatenation of the sensing matrixes corresponding to two different focal planes, and the recovery results are compared in Figure 6(c). We can see that although the SNR is reduced in a biplane observation, its recovery precision (as shown with our simulations) was higher due to the added constraints afforded by the biplane observation.

**Figure 6.**
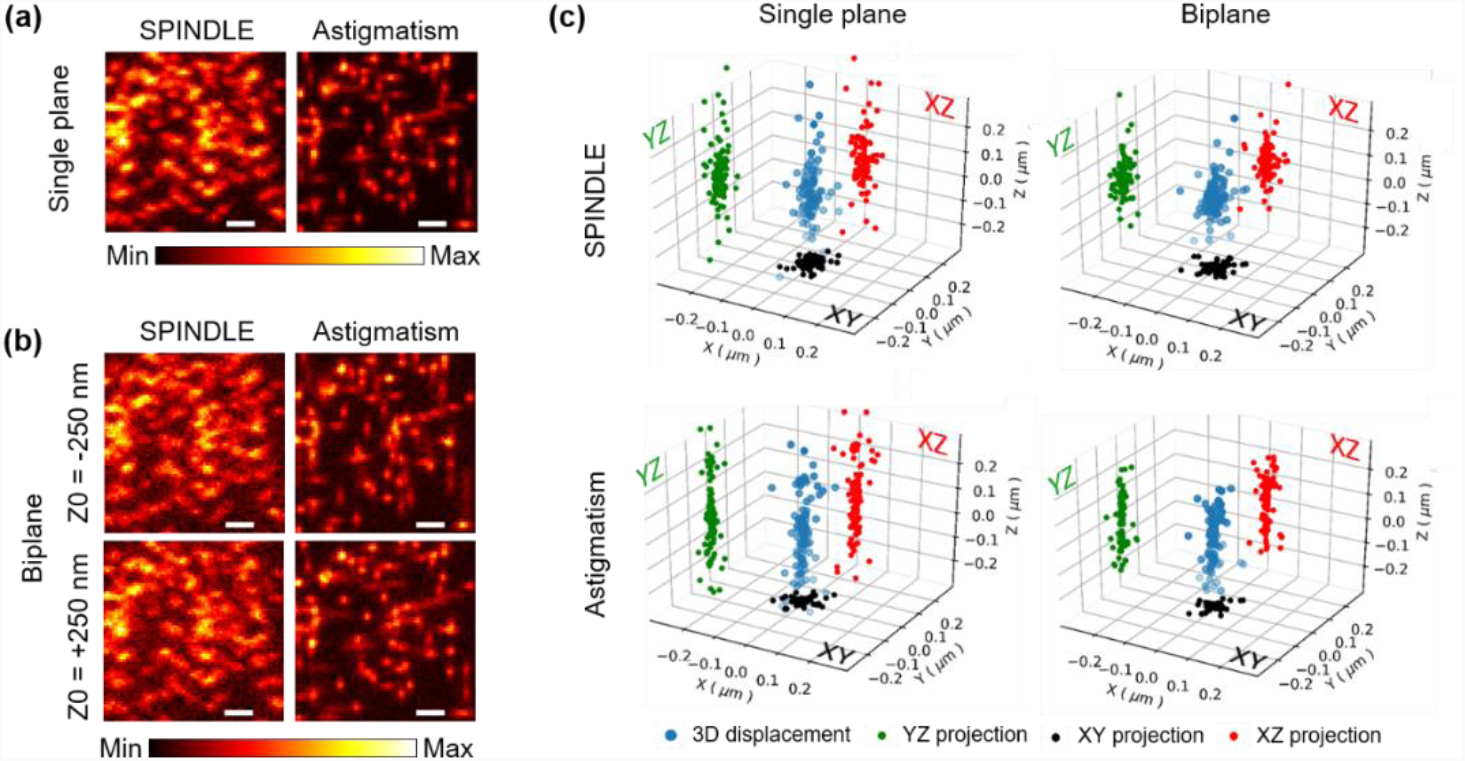
Validation of generalized PRIS recovery with vertical concatenation. PRIS recovery for biplane observation is demonstrated and compared to single-plane. The identical simulated sample is used for both observation configurations, using either SPINDLE PSF or Elliptical (astigmatic) PSF. (a) shows the input images for both PSF models using single plane observation and (b) shows the corresponding biplane observations, where the two focal planes are 125 nm apart from the focal plane used in single plane simulations. The recovery results are compared in (c), where each recovered localization coordinate is compared to the ground truth coordinate, and the displacement in three dimensions are plotted in the 3D scatter plot (with projections as shown in the graph). Four scattered plots demonstrate four different conditions with either single or biplane observations, using either SPINDLE or Astigmatic PSF. We can see that biplane improves localization precision for both PSF models, and Astigmatic demonstrate better precision in the lateral dimensions, while SPINDLE demonstrate better precision in the axial dimension. Scale bars: 1 μm.

### 3.3. Recovery with single-channel observation and dual-species (horizontal concatenation)

We further tested the generalization strategy of PRIS with the horizontal concatenation of the sensing matrixes (Figure 7). The simulation contains a mixture of two Cy3 molecules (577 nm emission) and two Cy5 molecules (690 nm emission) both detected through the same channel. The simulated detection scenario could be (but not limited to) dual laser excitation with a multi-band pass filter that allows for detection of both colors. For the detection path, we simulated a transparent phase mask that alters only the phase of the wavefront without phase wrapping in the fabrication. We set the phase mask to be a SPINDLE phase mask that matches 577 nm emission. Emission of both colors pass through the same focusing lens and phase mask, but the phases of the wavefront were altered differently by the phase mask and the defocus depth, due to different wavelengths, yielding different PSF profiles for each color. As shown in Figure 7(a), the PSF for Cy3 exhibit the expected SPINDLE PSF profile. However, for Cy5 molecule with mismatched emission wavelength used in the phase mask design, the PSFs exhibit extra central lobe and diffraction patterns (Figure 7(a)) that mismatches with a theoretical SPINDEL PSF profile. Such differences in the PSFs enable PRIS to distinguish two species by constructing the sensing matrixes A1 and A2 through the two PSFs separately as shown in Figure 7(b). The related experimental scenario could be independent characterizations of experimental PSFs with fluorescence beads of different colors through the same detection path.

In the common dual-color imaging system, either two filters are applied sequentially with sequential camera exposure, or the channel is split into two with different color filters. Such hardware approaches to separate color channels come at the cost of detected photon budget. This example demonstrates the separation of different color species mathematically by constructing the different species into the sensing matrix while capturing photons from both species in the same channel at the same time.

**Figure 7.**
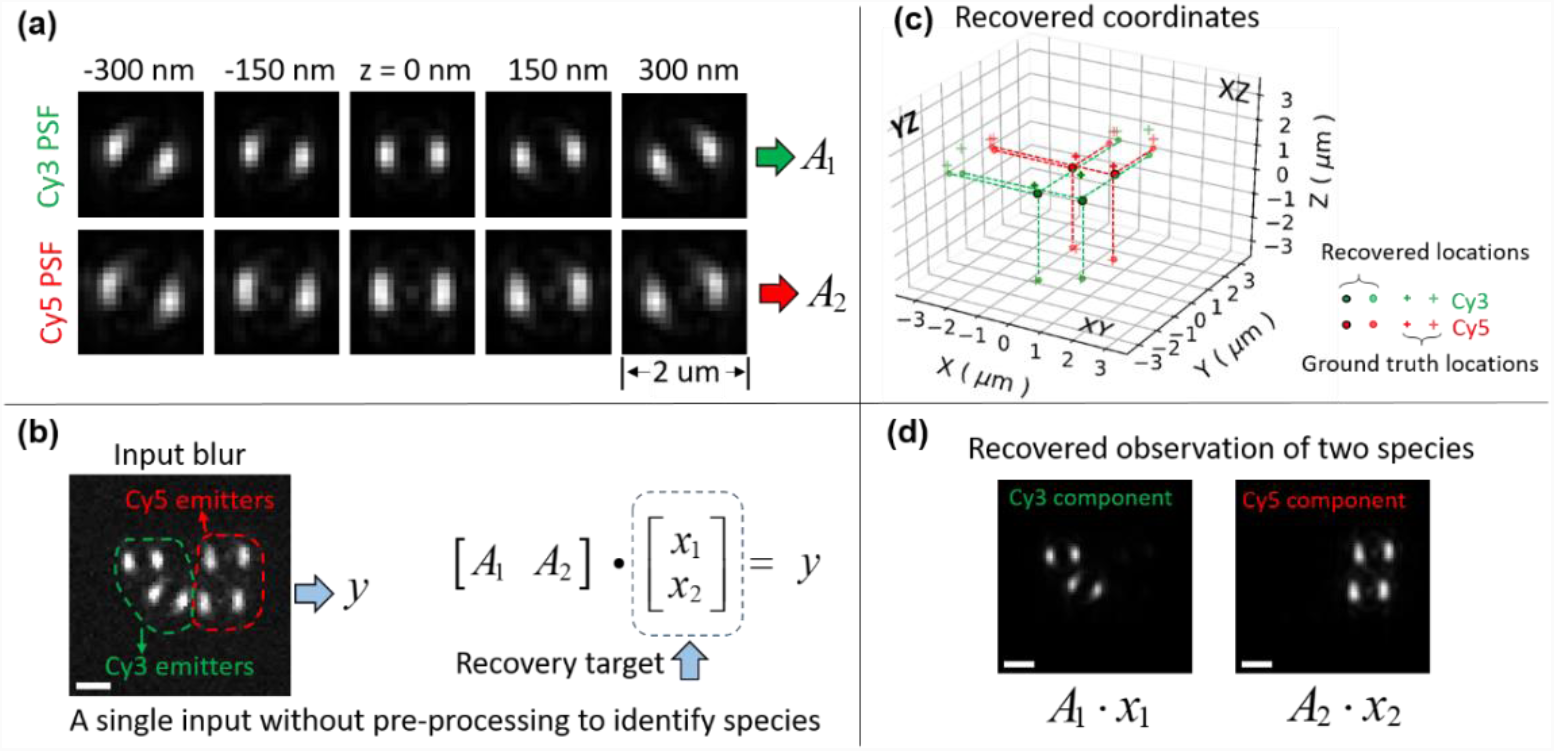
Validation of generalized PRIS recovery using horizontal concatenation. A single channel observation is simulated with a mixture of 4 fluorophores (two Cy3 molecules with 577 nm emission, and two Cy5 molecules with 690 nm emission). In the single observation channel, the SPINDLE phase mask is simulated to match the wavelength of Cy3. (a) shows the simulated 3D PSF slices for both fluorophores under the same detection path, for which Cy3 PSF shows the SPINDLE PSF profile, and Cy5 PSF shows extra diffraction lobes/patterns that would be unwanted in a traditional sense of double helix PSF application. (b) shows the simulated blurry noisy observation with the four emitters. The observed images of Cy3/Cy5 molecules are circled with green/red dashed lines respectively. For the sparse recovery problem, the single input image is vectorized to construct the ***y*** vector, two sensing matrixes constructed from the two species are horizontally concatenated to from the overall sensing matrix, and the sparse recovery target is the combined vector of ***x***_***1***_ and ***x***_***2***_. PRIS refinement are applied to two species separately. The final PRIS result is the recovery coordinates shown in (c) with species tag, and compared to the ground truth coordinates. (d) shows the observation reconstructed from the recovered ***x***_***1***_, ***y***_***2***_ with PRIS iterations. which clearly indicates the separation of two color species. Scale bars: 1 μm.

### 3.4. Recovery of curve feature with ultra-high labeling density

We further tested the recovery capability with a densely labeled curve in 3D with SPINDLE PSF (Figure 7(a)). The virtual sample is a curve with a total length of 8 μm, that expands approximately 5 μm in the lateral dimensions, and 0.56 μm in the axial dimension. A total number of 400 emitters are randomly placed along the line with cross-section location uncertainty of 7 nm, resulting in an average labeling density of 1 emitter per 20 nm. The emitters are all set at bright state in the simulation with randomly selected photon budgets between 4000 to 5000 and the background photon count is set to be 100 per pixel, and the photon counting noise is simulated with Poisson statistics. Figure 7(d) shows the simulated image measurement with the SPINDLE PSF. We note here that because SPINDLE PSF has two lobes with changing orientations to encode the defocus depth, the observation of the densely labeled curve in 3D appears as two lines. Figure 7(b)(c)(e) demonstrate the sparse recovery result of this simulated condition with different projection and zoomed panels as labeled in the figure. The sparse recovery is good in terms of recover the ground truth 3D curve, but without exhibiting individual localization coordinates. Additionally, because the intensity along the recovered line is rather smooth, the subsequent classification method couldn’t identify the coordinates of individual localization results. We attribute this to the ultra-high labeling density where the inter space between adjacent emitters are limited, therefore the reconstruction lacks the ability to distinguish individual emitters at this regime of high local density of emitter. However, as shown in Figure 7(b)(c)(e), the recovery result faithfully represents the ground truth feature as a curve.

**Figure 7.**
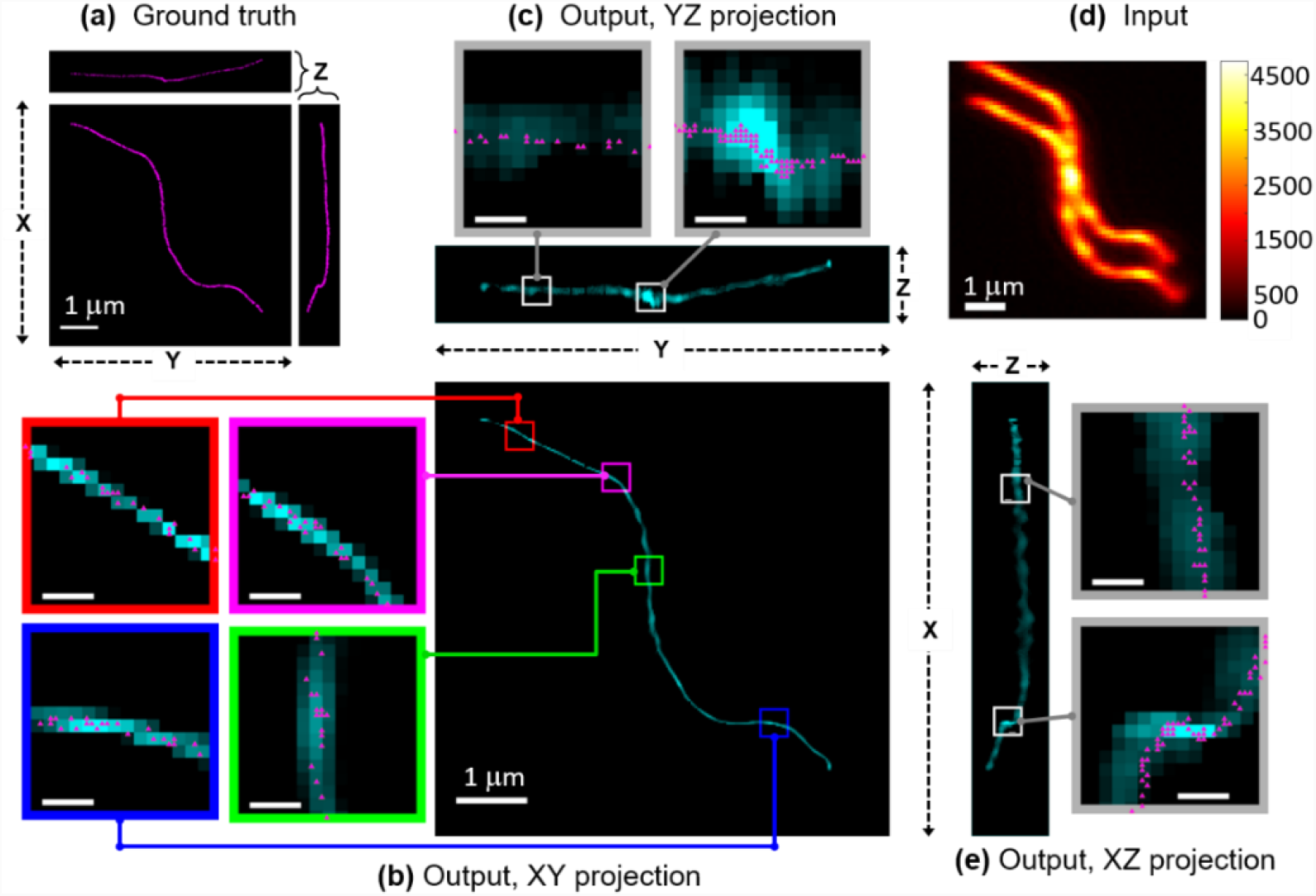
PRIS recovery of densely labeled curve in 3D. PRIS result of a simulate 3D densely labeled (1 emitter per 20 nm on average) curve is shown here. (a) shows the ground truth emitter locations in projection of XY-plane (center), XZ-plane (right) and YZ-plane (top). Each emitter location is labeled as a magenta dot. The displays of the ground truth appear as curves due to the high-density of labeled emitters. The simulated measurement with SPINDLE PSF is shown in (d). Note that because of the nature of the double-helix PSF (two lobes), the image of a single curve appears as two curves in the observation. The sparse recovery results are shown in (b), (c) and (e) with different projection views with zoomed panels as labeled in the major panel and connected to the zoomed panels. In the zoomed panels, the ground truth emitters are marked with magenta triangles. We can see that although the sparse recovery does not recover the location coordinates of individual emitters, but the result represents the underlying 3D curve as compared to the ground truth. Scale bars in major panels are 1 as labeled in the figure; scale bars in zoomed panels without labels are 100 nm.

## 4. CONCLUSIONS AND DISCUSSION

In this work, we developed a progressive refinement method for compressive sensing (PRIS) to perform 3D super-resolution recovery of fluorescence microscopy observations, without reliance on the convolution assumption of the imaging process. The method is generalized to work with different PSF models, multiple observation channels and a mixture of species of the signal sources. We have demonstrated high-fidelity reconstructions of simulated data sets for double-helix PSF (SPINDLE), astigmatic PSF, and for both single and biplane image acquisitions. We also demonstrated the recovery capability with a densely labeled curve in 3D. Our work is useful for the algorithmic synchronization of different imaging modalities, where the synchronization is realized at the stages of data acquisition and data reconstruction. It affords a deeper level of synchronization as compared to sequential data analysis. In the more general sense, PRIS can be generalized to sparse recovery even at a hyperdimensional space with tensor formulation[30], and the systematic progressive refinement can be applied at each dimension (each mode of the tensor) independently and/or collectively. Our method can also be applied to observations with well isolated PSFs in the observation image for single molecule localization microscopy. Accordingly, proper initialization of PRIS iteration can be used with voxels covering only identified interested areas to reduce the computation cost further. Least square fittings can be applied on top of our algorithm to enhance the localization precision further. The highly generalized form and computation simplicity allow for scalable recovery platforms, such as cloud computing, GPU, and FPGA implementations.

## ACKNOWLEDGEMENT

The first author would like to thank Prof. Jing Qin, Prof. Wotao Yin, Prof. Bo Huang, Prof. Giang Tran, Prof. Stanley Osher, Dr. Yi Xia, Mr. Minh Pham and Ms. Wenbo Gu for the inspirational discussions concerning theory and applications of L1-norm regularized sparse recovery and compressive sensing. This work is supported by NSF STROBE: A National Science Foundation Science & Technology Center under Grant No. DMR 1548924. This work used computational and storage services associated with the Hoffman2 Shared Cluster provided by UCLA Institute for Digital Research and Education’s Research Technology Group.

## APPENDIX

**Table 1.**
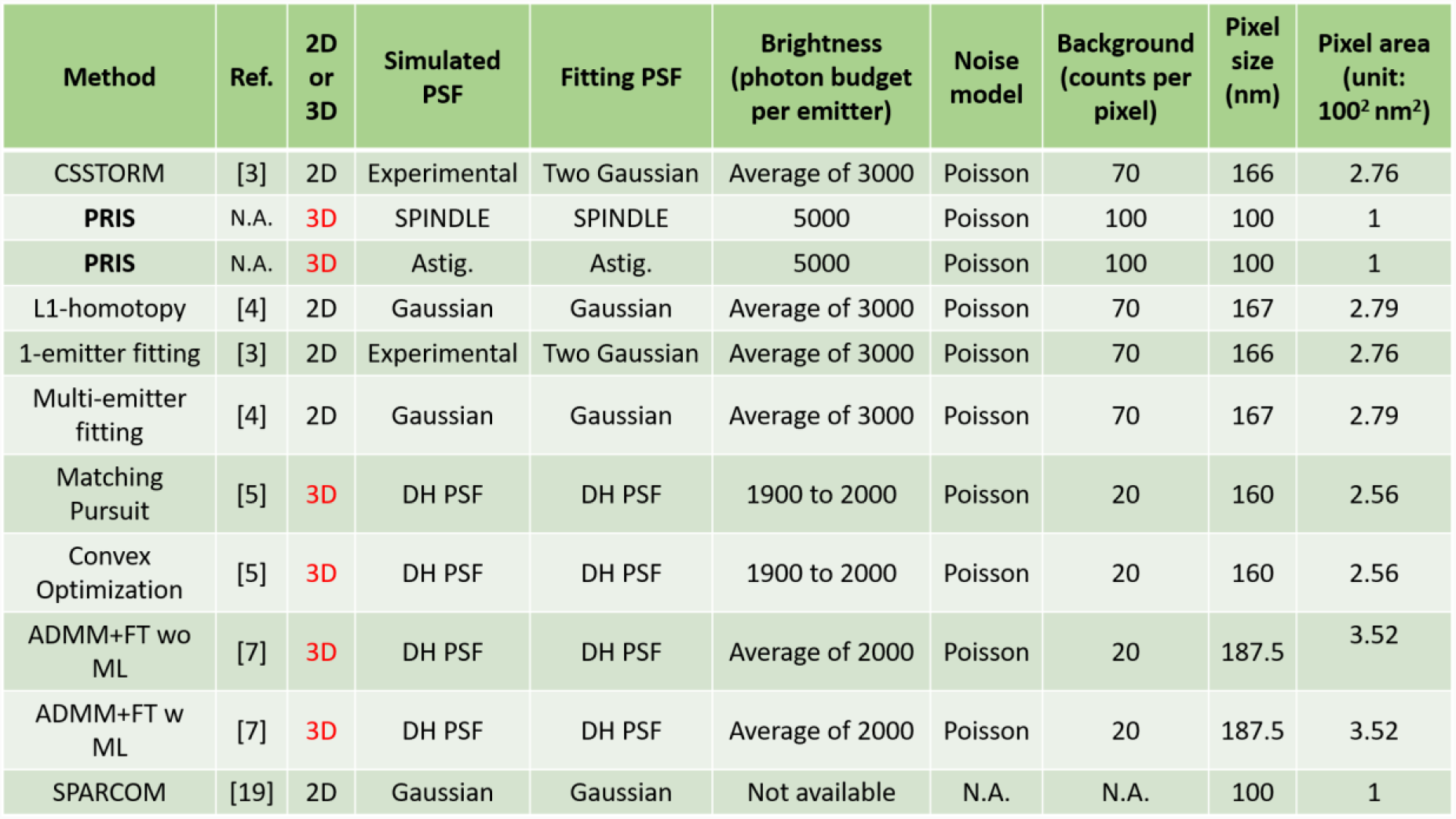
Simulation conditions used for the recovery performance quantification in previous works[3-5, 7, 20] and plotted in Figure 5.

